# High-resolution eye-tracking via digital imaging of Purkinje reflections

**DOI:** 10.1101/2022.08.16.504076

**Authors:** Ruei-Jr Wu, Ashley Clark, Michele Cox, Janis Intoy, Paul Jolly, Zhetuo Zhao, Michele Rucci

## Abstract

Reliably measuring eye movements and determining where the observer looks are fundamental needs in vision science. A classical approach to achieve high-resolution oculomotor measurements is the so-called Dual-Purkinje-Image (DPI) method, a technique that relies on the relative motion of the reflections generated by two distinct surfaces in the eye, the cornea and the back of the lens. This technique has been traditionally implemented in fragile and difficult to operate analog devices, which have remained exclusive use of specialized oculomotor laboratories. Here we describe progress on the development of a digital DPI, a system that builds on recent advances in digital imaging to enable fast, highly precise eye-tracking without the complications of previous analog devices. This system integrates an optical setup with no moving components with a digital imaging module and dedicated software on a fast processing unit. Data from both artificial and human eyes demonstrate sub-arcminute resolution at 1 Khz. Furthermore, when coupled with previously developed gaze-contingent calibration methods, this system enables localization of the line of sight within a few arcminutes.

## Introduction

Measuring eye movements is a cornerstone of much vision research. While the human visual system can monitor a large field, examination of fine spatial detail is restricted to a tiny portion of the retina, the foveola, a region that only covers ~1° in visual angle, approximately the size of a dime at arm-length. Thus, humans need to continually move their eyes to explore visual scenes: rapid gaze shifts, known as saccades, redirect the foveola every few hundred milliseconds, and smooth pursuit movements keep objects of interest centered on the retina when they move. Oculomotor activity, however, is not restricted to this scale. Surprisingly, eye movements continue to occur even during the so-called periods of fixation, when a stimulus is already examined with the foveola. During fixation, saccades that are just a fraction of a degree (microsaccades), alternate with an otherwise incessant jittery movement (ocular drift) that yields seemingly erratic motion trajectories on the retina. Much research has focused on the characteristics and visual functions of these movements^1–3^.

Thus, eye movements vary tremendously in their characteristics, ranging by over two orders of magnitude in amplitude and three orders of magnitude in speed, from a few arcmimutes/s to hundreds of degrees/s. This wide range of motion, together with the growing need to resolve very small movements, pose serious challenges to eye-tracking systems. Although various methods for measuring eye movements have been developed, several factors limit their range of optimal performance. Such factors include changes in the features that are tracked (*e.g.* the pupil, the iris), translations of the eye as those caused by head movements, and uncertainty in converting raw measurements into gaze positions in the scene.

A technique that tends to be particularly robust to artifacts and combines high resolution with fast dynamics has long been in use, but it has remained confined to specialized oculomotor laboratories. This method, known as Dual Purkinje Imaging (DPI), relies on tracking cues from more than one surface in the eye, rather than just the cornea as done by most eyetrackers (Fig. 1A). Specifically, it compares the positions of reflections of an infrared beam of light from the anterior surface of the cornea (the 1st Purkinje image) and the posterior surface of the lens (the 4th Purkinje image; Fig. 1B). These two images move relative to each other as the eye rotates, and their relative motion is little affected by eye translations. Thus, the DPI method enables oculomotor measurements with high spatial and temporal resolution, while offering robustness to artifacts caused by head movements and changes in pupil size.

**Figure 1:**
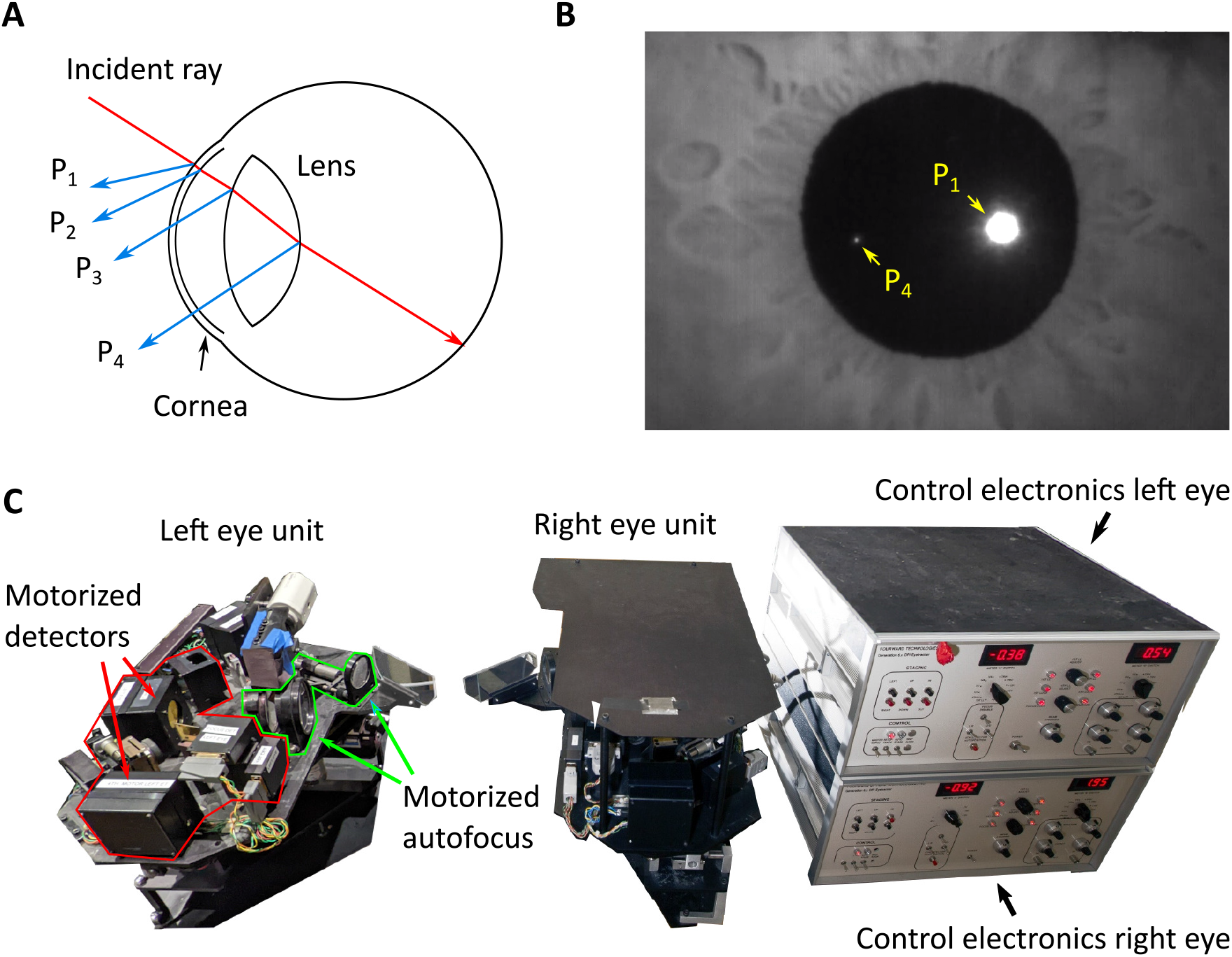
Purkinje reflections and eye tracking. (**A**) As it travels through the eye, an incident ray of light gets reflected by the structures that it encounters. The first and second Purkinje reflections (P_1_ and P_2_) result from the anterior and posterior surfaces of the cornea. The third and the fourth Purkinje reflections (P_3_ and P_4_) originate from the anterior and posterior surface of the lens. (**B**) An example of Purkinje images. P_1_ and P_4_ can be imaged simultaneously. Here P_3_ is out of focus, and P_2_ largely overlaps with P_1_. Note that P_1_ is larger and has higher intensity than P_4_. (**C**) An example of eye-tracker that uses Purkinje reflections, the Generation V Dual Purkinje Eye-tracker (DPI) from Fourward Technologies. This optoelectronic device has mobile components that, when properly tuned, follow P_1_ and P_4_ as the eye rotates.

Furthermore, this method can deliver measurements with minimal delay, paving the way for procedures of gaze-contingent display control—the updating of the stimulus in real-time according to eye movements^4^. This is important not just for experimental manipulations, but also because gaze-contingent calibrations allow translating precision into accuracy, *i.e.* they lead to better determination of where the observer looks in the scene^5^. This approach has been used extensively to investigate the purpose of small saccades^6,7^, the characteristics of foveal vision^8,9^, and the dynamics of vision and attention within the fovea^10,11^.

So far, the DPI principle has been exclusively implemented in analog opto-electronic devices. These systems, originally conceived by Cornsweet and Crane^12,13^, use servo feedback provided by analog electronics to control the motion of detectors so that they remain centered on the two Purkinje images (Fig. 1C). Although capable of resolving eye movements with arcminute resolution, several features of DPI eye-trackers have prevented their widespread use. A major limiting factor has been the difficulty to both operate and maintain the device. Sophisticated calibration procedures are required for establishing the proper alignment in the optical elements, preserving their correct motion, and also tuning the system to individual eye characteristics. Since several components move during eye-tracking, these devices are fragile and suffer from wear and stress, requiring frequent maintenance. As a consequence, few DPI eye-trackers are currently in use, most of them in specialized oculomotor laboratories.

The recent technological advances in CCD cameras, image acquisition boards, and computational power of digital electronics now enable implementation of the DPI approach in a much more robust and easily accessible device. Here we report recent progress in the development of a digital DPI eye-tracker (the dDPI), a system that uses a fast camera, high-speed image analysis on a dedicated processing unit, and a simple but robust tracking algorithm to yield oculomotor measurements with precision and accuracy comparable to those of the original analog device. In the following sections, we first review the principles underlying DPI eye-tracking. We then describe a prototype dDPI and examine its performance with both artificial and real human eyes.

### Measuring eye movements from Purkinje reflections

As shown in Fig. 1A, the Purkinje images are the reflections given by the various eye structures encountered by light as it travels toward the retina. Because of the shape of the cornea and lens, the first and fourth Purkinje images, *P*_1_ and *P*_4_, form on nearby surfaces and can be imaged together without the need of complex optics. Since these two reflections originate from interfaces at different spatial locations and with different shapes, they do not move together as the eye rotates. Thus, the relative motion between *P*_4_ and *P*_1_ provides information about eye rotation. Below we examine the main factors that need to be taken into consideration for using this signal to measure eye movements.

As originally noted by Cornsweet and Crane (1973),the relative motion of *P*_1_ and *P*_4_ follows a monotonic relation with eye rotation. An approximation of this relation can be obtained, for small eye movements, by using a paraxial model. In Fig. 2A, the anterior surface of the cornea and the posterior surface of the lens are replaced by a convex and a concave mirror (*M_C_* and *M_L_*), respectively. Under the assumption of collimated illumination, this model enables analytical estimation of the relative positions, *X*_1_ and *X*_4_, of the two Purkinje images on the image plane *I*:

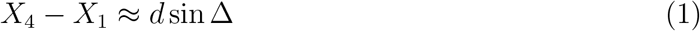

where *d* represents the distance between the anterior surface of cornea and the posterior surface of the lens; and Δ is the angle between the incident beam of light and the optic axis of the eye, which represents here the angle of eye rotation (see Fig. 2A).

**Figure 2:**
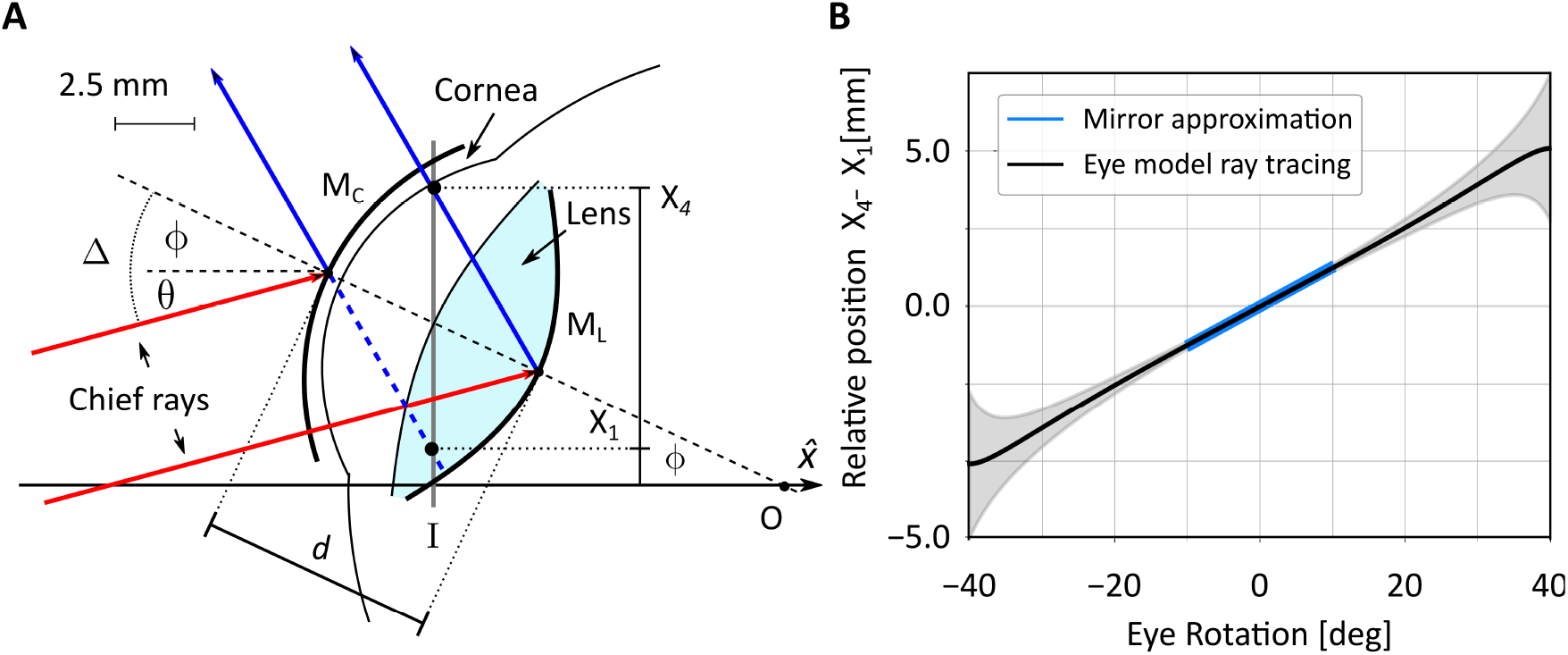
A paraxial model of Purkinje images formation. (**A**) Schematic diagram of the eye with the anterior surface of the cornea and the posterior surface of the lens approximated by two mirrors (*M_C_* and *M_L_*). The optical axis of the eye is initially aligned with that of the imaging system 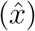 and is illuminated by a collimated beam tilted by *θ*. The eye then rotates by *ϕ* around its rotation center *O*. Light (here represented by the chief rays in red) is reflected by the cornea and the lens (blue lines) so that it intersects the image plane *I* at locations *X*_1_ and *X*_4_, respectively. These are the positions were the Purkinje reflections P_1_ and P_4_ form in the image. (**B**) Predicted relative motion of the two Purkinje reflections as the eye rotates. Both the motion predicted for small eye rotations by the paraxial mirror model in *A* (blue line) and the motion predicted over a broader range of eye rotations by a more sophisticated model (black line, see Fig. 3) are shown. Shaded regions represent standard errors obtained by varying eye model parameters to reflect normal individual variability. Incident light is here assumed parallel to the optical axis of the imaging system (*i.e., θ* = 0 in *A*).

In this study, we focus on the case of the unaccommodated eye. Inserting in Eq. 1 the value of d taken for Gullstrand eye model (*d* = 7.7 mm) and assuming light to be parallel to the optical axis of the imaging system (*θ* = 0 in Fig. 2A), we obtain the blue curve in Fig. 2B. This model suggests that, for the sufficiently small range of eye movements for which the paraxial model holds, the relative positions of the two Purkinje images varies almost linearly with eye rotation.

While the two-mirror approximation of Fig. 2A is intuitive, it holds only for small eye movements, *i.e.,* for small angular deviations between the illumination axis and the line of sight. Furthermore, it does not consider the presence of other refracting structures in the eye, nor the non-homogeneous optical characteristics of the human lens. To gain a deeper understanding of the positions and shapes of the Purkinje images and how they move across a wide range of eye rotations, we performed ray-tracing of a more sophisticated eye model. A primary goal of these simulations was to determine the expected sizes, shapes, and strengths of the Purkinje images as the angle of incident light changes because of eye rotations. Being reflections, these characteristics depend both on the source of illumination and the optical/geometrical properties of the eye.

The model used here is based on multiple anatomical and optical measurements from human eyes. It has been previously validated in several ways and compared to other schematic eye models^14,15^. This model consists of 6 refracting surfaces and a lens with a gradient refractive index designed to model the spatial non-homogeneity present in the human lens ^16^. It was simulated in Zemax OpticStudio with parameters adjusted to model the unaccommodated emmetropic eye with a pupil of 6 mm. It was exposed to a collimated uniform beam with a 10 mm diameter, a beam sufficiently wide to cover the pupil under the entire range of considered rotations.

Since our goal is to develop a system that can be used in vision research experiments, we illuminated the model with light in the infrared range (850 nm), so that it would not interfere with visual stimulation. Under exposure to collimated light, eye rotations only change the angle of incident light relative to the optical axis of the eye. We, therefore, modeled eye movements by keeping the model stationary and varying the illumination angle relative to the eye. In all cases, we modeled Purkinje images formation on the plane where *P*_4_ gives on average maximum image irradiance (*I* in Fig. 2A).

We first examined the impact of the power of the illumination on Purkinje image formation. Fig. 3A and B show one of the primary difficulties of using Purkinje images for eye-tracking: the irradiance of *P*_1_ and *P*_4_ differs considerably, by almost two order of magnitudes. As expected, both vary proportionally with the power of the source and decrease with increasing angle between the optical axis of the eye and the illuminating beam. However, their ratio remains constant as the power of the source varies, and the vast difference in irradiance between the two makes it impossible—in the absence of additional provisions—to image *P*_4_ without saturating *P*_1_.

**Figure 3:**
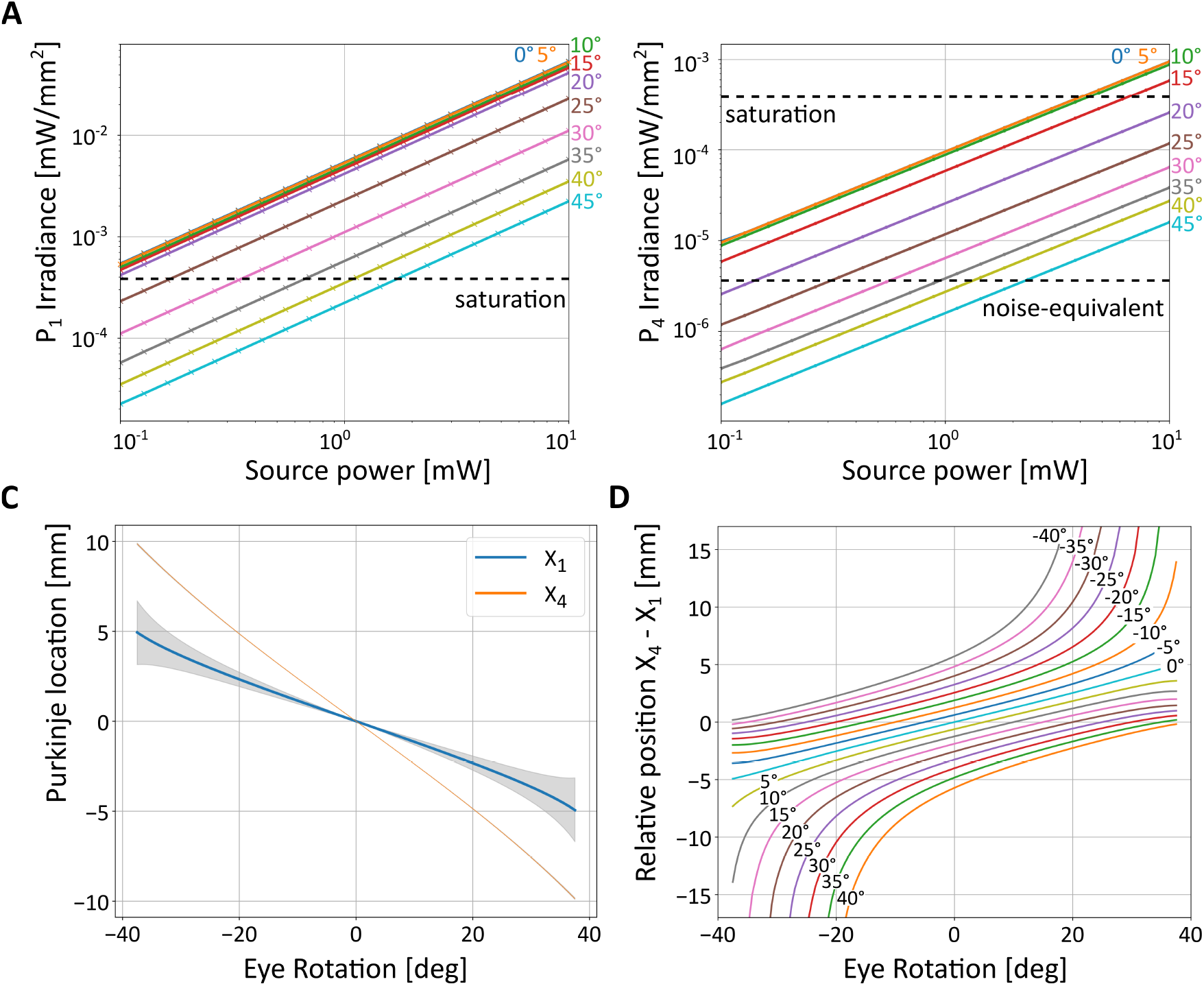
Expected characteristics of Purkinje images from ray-tracing a model of the eye. (**A-B**) Maximum irradiance of the two Purkinje images as a function of the power of the illumination source. Each curve shows data for a specific angle of eye rotation, measured as deviation between the optical axes of the eye and the imaging system (the latter assumed to be coincident with the illumination axis, *θ* = 0 as in Fig. 2*B*). The two horizontal dashed lines represent the minimum and maximum intensity measurable with the sensor in our prototype. (**C**) Locations of the peak intensity of the two Purkinke images as a function of eye rotation. Shaded regions (not visible for P4) represent standard errors obtained by varying eye model parameters to reflect normal individual variability. (**D**) Relative position of *P*_1_ and *P*_4_ as a function of both eye rotation and illumination angle.

These data are informative for selecting the power of the illumination. There are two primary constraints that need to be satisfied: from one side, the power of the source needs to be sufficiently low to meet safety standards for prolonged use. From the other, it needs to be sufficiently high to enable reliable detection of the Purkinje images. The latter requirement is primarily determined by the noise level of the camera, which sets a lower bound for the irradiance of *P*_4_, the weaker reflection. Fig. 3A and B show the saturation and noise-equivalent levels of irradiance for the camera used in our prototype (see Section 3). A power source higher than 1 mW yields a detectable *P*_4_ (Fig. 3B), but the eye-tracking algorithm needs to be able to cope with the simultaneous saturation of large portions of *P*_1_ (Fig. 3A). This model also allows more rigorous simulation of how the two Purkinje images will move in the image than the paraxial approximation of Fig. 2A. Fig. 3C shows the chief rays intersections with the image plane, *X*_1_ and *X*_4_, for eye rotations of ± 40 degrees. Both images are displaced almost linearly as a function of eye rotation. However, the factor of proportionality differs between *P*_1_ and P4, with the latter moving substantially faster and, thus, covering a broader extent. These data indicate that the imaging system should capture approximately 20 mm to enable measurement of eye movements over this range. The model also shows that motion characteristics are minimally influenced by individual changes in eye morphology and optical properties, as captured by the normal range of model parameters measured in healthy eyes^16^ (shaded regions in Fig. 3C).

Since both Purkinje images move proportionally to eye rotation, also the difference in their positions maintains an approximately linear relation with eye movements. This function closely matches the paraxial two-mirror approximation of Eq. 1 for small eye movements and extends the prediction to a much wider range of eye rotations (the black line in Fig. 2B). In the simulation of this figure, the direction of illumination and the optical axes of both the eye and the imaging system are all aligned. However this configuration is not optimal, as the two Purkinje images overlap when the optic axis of the eye is aligned with the illumination axis. This makes it impossible to detect *P*_4_, which is much smaller and weaker than *P*_1_. As shown in Figure 3D, the specific position of the illuminating beam plays an important role, both in determining departures from linearity and in placing the region where *P*_4_ will be obscured by *P*_1_. In this figure, the offset between the two Purkinje images is plotted for a range of eye rotations (x-axis) and for various angular positions of the source (distinct curves). In the system prototype described in this article (Fig. 5), we placed the source at *ϕ* = 20° on both the horizontal and vertical axes. This selection provides a good linear operating range, while placing the region in which *P*_4_ is not visible away from the most common directions of gaze.

## A fast tracking algorithm

Once the Purkinje images are acquired into a digital format, a variety of methods are available for their localization and tracking. In the traditional analog DPI device, the system dynamics is primarily determined by the inertia of the mobile components and the filtering of the servo signals, which yield a broad temporal bandwidth^13^. As a consequence, analog DPI data are normally sampled at 500 Hz-1 kHz following analog filtering at the Nyquist frequency^17–20^. In this system, the overall delay with which data are made available is sufficiently small to enable procedures of gaze-contingent display control, experiments in which the stimulus is modified in real-time according to the eye movements performed by the observer. Here we describe a simple tracking algorithm that mimics the processing of the analog DPI and, when implemented in a fast processor like a Graphic Processing Unit (GPU), is capable of delivering measurements at 1 kHz with delay smaller than 1 ms. m The design of an efficient tracking algorithm needs to take into consideration the different characteristics of the two Purkinje images. Fig. 4A shows the images predicted by the model eye of Section 2 when a low-power (2 mW) source illuminates the eye from 20° off the camera’s axis both horizontally and vertically, as in the prototype of Fig. 5. Two observations immediately emerge. First, the two reflections differ considerably in size, as *P*_4_ has much lower intensity that *P*_1_ and will appear in the image as a small blob compared to the first reflection. Second, as pointed out in the previous section, direct imaging of the eye will yield an image of *P*_1_ that is largely saturated and an image of *P*_4_ close to the noise level. Thus, the localization algorithm should be both tolerant to saturation effects and insensitive to noise.

**Figure 4:**
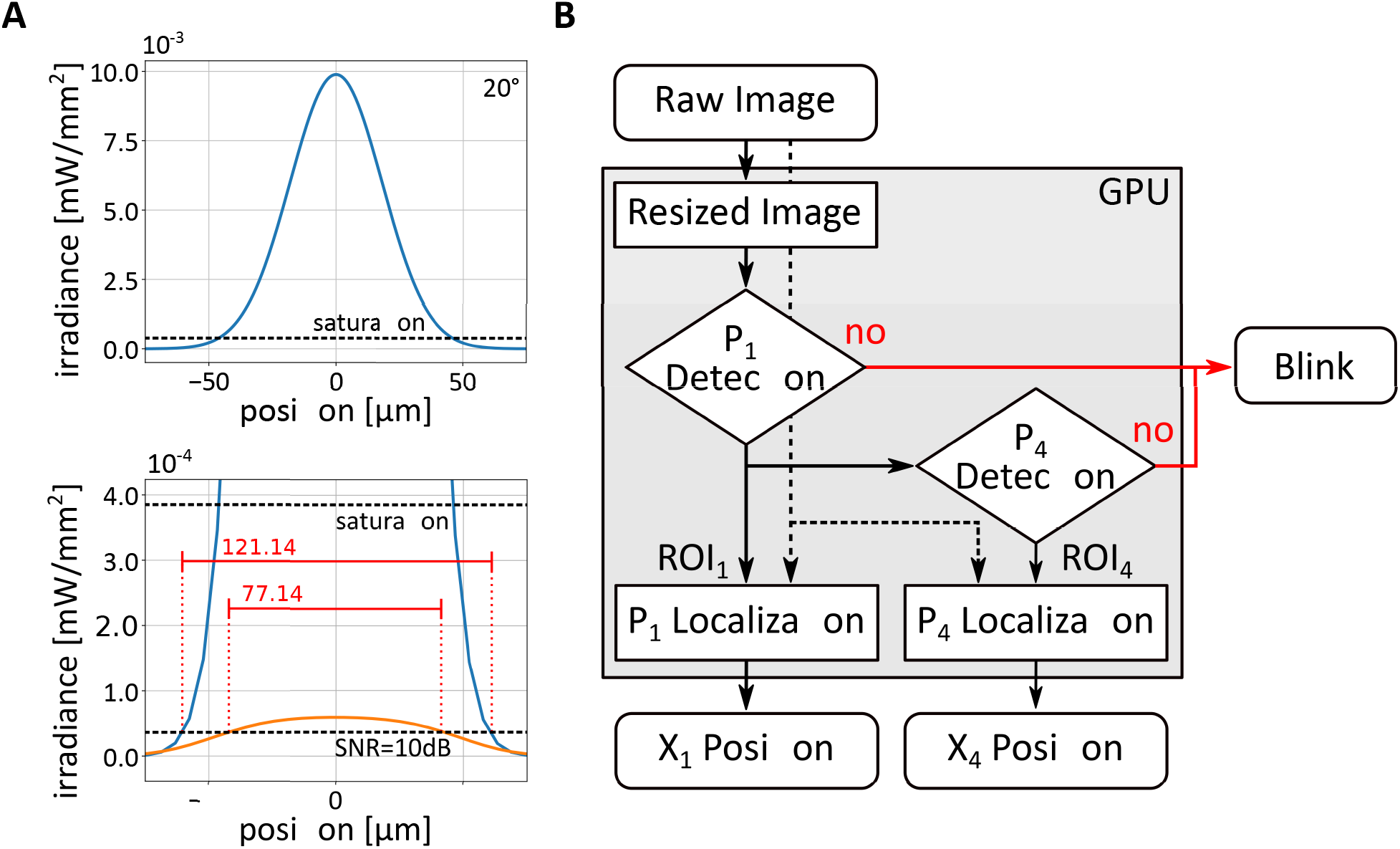
A fast tracking algorithm. (**A**) Characteristics of Purkinje images. Data are obtained from the model in Fig. 3, using a source power of 2 mW located at 20° on both axes. The top panel shows the profile of *P*_1_ before clipping intensity at the saturation level. The bottom panel shows both *P*_1_ (with saturated intensity; blue curve) and *P*_4_ (orange curve). The red horizontal bars represent the diameters of the reflections exceeding 10 dB. (**B**) Flowchart of the tracking algorithm. A low-resolution image of the eye (solid line) is first used to detect P_1_ and P_4_ and determine their approximate locations. Failure in detecting one of the two reflections is labeled as blink. High-resolution localization of the Purkinje images (dashed lines) is then obtained from selected region of interests (ROI_1_ and ROI_4_).

**Figure 5:**
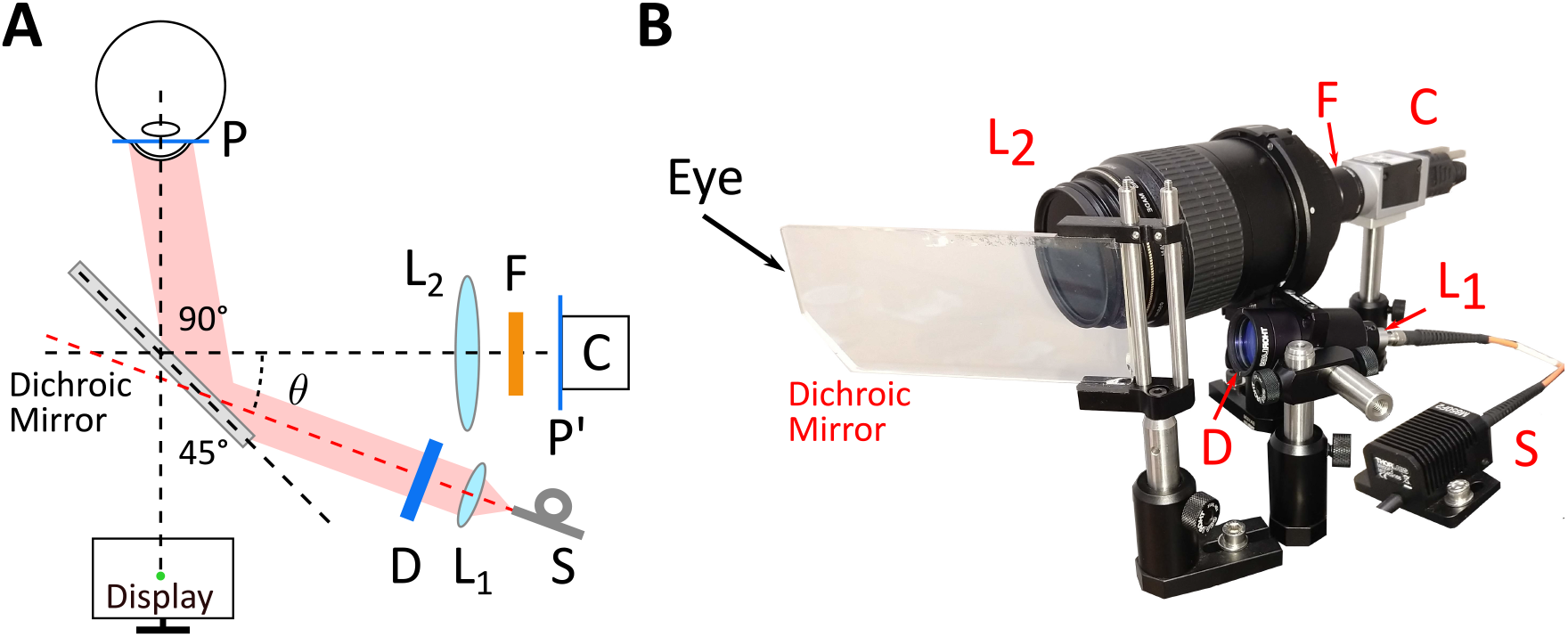
A dDPI prototype. (**A**) General architecture and (**B**) hardware implementation. Infrared light from the source (*S*, a fiber-coupled LED; Thorlabs M850F2 and M28L01W) is collimated (lens *L*_1_; Thorlabs F260SMA-850) and expanded to fully cover the eye by a beam expander (*D*; 3X). The beam is directed toward the eye via a dichroic mirror. Purkinje reflections are acquired by a high-speed camera (*C*; IO Industries Flare 12M180CX) equipped with a long-pass filter (*F*; Thorlabs FEL0800). The camera lens (*L*_2_; Canon EF 100mm f/2.8L) relays the plane of the Purkinje images *P* to the sensor plane *P*′.

The algorithm used in this study is shown in Fig. 4B. It operates directly on the digital images of the two Purkinjes and yields as output the differences between the Purkinje positions on each of the two Cartesian axes of the image. To ensure low latency, the algorithm operates in two phases, detection and localization. In the first phase, it defines regions of interest (ROIs) in which *P*_1_ and *P*_4_ are highly likely to be found. In the localization phase, it then operates on the selected regions at high resolution to precisely determine the coordinates of *P*_1_ and *P*_4_.

The detection phase operates on downsampled versions of the acquired images, which are for this purpose decimated by factor 8. Different approaches are used for the two reflections, given their distinct characteristics in the images. Since *P*_1_ is relatively large and typically the only saturated object in the image, its region of interest (ROI_1_, a 256 × 256 pixel square) can be directly estimated by computing the center of mass of a binary thresholded image, as long as a sufficiently high threshold is used to avoid influences from other possible reflections in the image. To further speed up computations, this approach can be applied to a restricted portion of the input image, since *P*_1_ moves less that *P*_4_ and remains within a confined region of the image.

Detection of *P*_4_ is more complicated given its much lower intensity and smaller size. The algorithm in Fig. 4B uses template matching, an operation that can be executed rapidly using the parallel hardware of modern GPUs. Specifically, the algorithm estimates the location in the image that best matches—in the least squares sense—a two-dimensional Gaussian template of *P*_4_, the parameters of which had been previously calibrated for the individual eye under observation. This yields a 64×64 pixel square region centered around *P*_4_ (ROI_4_). Like for the computation of ROI_1_, if needed, also this operation can be restricted to a selected portion of the image to further speed up processing, in this case using knowledge from *P*_1_ to predict the approximate position of *P*_4_.

In the second phase of the algorithm, ROI1 and ROI4, now sampled at high resolution, are used for fine localization of the two reflections. To this end, the position of each Purkinje was marked by its center of symmetry (the radial symmetry center), a robust feature that can be reliably estimated via a non-iterative algorithm developed for accurate tracking of particles^21^. This algorithm enables subpixel resolution, it is robust to saturation effects and has fast execution time.

This is shown in Table 1, which compares three different methods used for estimating the position of *P*_4_ in images obtained from ray-tracing the model described in Section 2. In these images the exact position of *P*_4_ is known, providing ground truth to compare approaches. As shown by these data, Gaussian fitting via maximum likelihood gives the highest accuracy, but this method is computationally expensive and cannot be executed in real time. In contrast, estimation of the center of mass can be performed very rapidly, but it yields a much larger error. Estimation of the radial symmetry center provides an excellent trade-off between accuracy and speed, with error comparable to Gaussian fitting, but computation time lower than 0.5 ms when implemented in a good quality GPU.

**Table 1:** Performance of three methods for localization of Purkinje reflections.

| Method | RMS (arcmin) | time by CPU (ms) | time by GPU (ms) |
| --- | --- | --- | --- |
| Gaussian Fitting | 0.021 | 3327 | - |
| Center of Mass | 0.22 | 34 | 0.05 |
| Radial Symmetric Center | 0.06 | 521 | 0.32 |

## System prototype

The data obtained from the model eye and simulations of the tracking algorithm enabled design and implementation of a 1-Khz prototype. A major requirement in the development of this apparatus was that it could be successfully used in vision research experiments in place of its analog counterpart; that is, that the device would be capable of fully replacing the traditional analog DPI in all relevant functional aspects. The system shown in Fig. 5 is based on a simple optical/geometrical arrangement and consists of just a few important components: a camera, an illuminator, a dichroic mirror, in addition to several accessory elements. Unlike its analogical predecessor, none of the components moves, guaranteeing robustness and limited maintenance.

The heart of the apparatus is a digital camera that acquires images of the Purkinje reflections. Selection of the camera impose a trade-off between spatial and temporal resolution, as higher sampling rates can only be achieved with smaller numbers of pixels. The high-speed camera in Fig. 5B (IO Industries Flare 12M180CX) was chosen based on our goal of obtaining 1′ spatial resolution and temporal resolution of ~ 1 ms. This camera has pixels that are just 5.5 *μ*m wide and delivers 187 frames per second at 12 MP. It can achieve 1 KHz acquisition rate at 2 MP.

Positioning of the camera was based on two factors. To reach the necessary magnification and be able to track over a wide range of eye movements, the camera was placed optically in front of the eye. To avoid obstructing the field of view and enabling clear viewing of the display in front of the subject, the camera was physically placed laterally to the eye, at a 90° angle from the eye’s optical axis at rest. This was achieved by means of a custom short-pass dichroic mirror that selectively reflected infrared light at a 45° angle while letting through visible light (see Fig. 5A).

Illumination is provided by a fiber-coupled LED at 850 nm with maximum power of 2 mW, well within the safety range for extended use. The beam is collimated and expanded by a factor of three to ensure that the eye remains uniformly illuminated over the relevant range of eye rotations. Based on the simulation data from Fig. 4, we placed the illuminator 20° off the optical axis of the camera on both the horizontal and vertical axes. This ensures that P1 and P4 will never overlap within the visual field covered by the monitor. This configuration also guarantees good linearity of the system in the range of considered eye movements.

Choice of the camera and its operating characteristics establishes a trade-off between spatial and temporal resolutions. This happens because of the hardwired limits in the camera’s transmission bandwidth, which forces the acquisition of larger images to occur at slower rates. In our prototype, based on our experimental needs, we prioritized high speed binocular tracking (1 Khz) at the expenses of the width of the field of view. Our simulations indicate that the first and fourth Purkinje images move relative to each other at the rate of 2.4 *μ*m per arcminute of eye rotation. Since the center of radial symmetry algorithm yields subpixel resolution provided that a sufficient number of pixels is available for each reflection, we aimed at covering *P*_4_—the smallest image—with more than 25× 25 pixels. This guarantees resolution higher than 1′ as shown in the simulations of Table 1 and confirmed below with controlled rotations of an artificial eye. These considerations led to the choice of an optical magnification of 1.25, with each pixel covering 3′.

By determining the field of view of the camera, optical magnification also defines the operating range of the eye-tracker. This is range in which *P*_4_ can be tracked, the reflection that moves the most as the eye rotates. Our simulations indicate that *P*_4_ moves by about 4.7 mm for 20° eye rotation. With the portion of our camera’s sensor that can be used at 1 Khz (5.632 mm; a quarter of the actual sensor), the resulting trackable range is about ±19°. Selection of different parameters would provide different trade-offs between spatial and temporal performance. For example, a much larger field can be tracked at high spatial resolution by acquiring data at 500 Hz.

The tracking algorithm shown in Fig. 4 was implemented in parallel processing (CUDA platform) in a state-of-the-art GPU (Nvidia Geforce 1080 Ti). This implementation enabled delivery of binocular data in less than a millisecond. The difference between the positions of the two Purkinje images given by the algorithm was converted in degrees of visual angles using a bilinear interpolation procedure estimated during a preliminary calibration. This procedure converts Purkinje image displacements into gaze direction and is identical to the one often performed in previous studies with the analog DPI device^5^.

## Results

To thoroughly characterize the performance of our prototype, we examined measurements obtained with both artificial and human eyes. Artificial eyes were used to provide ground truth baselines both in stationary tests and in the presence of precisely controlled rotations. Human eye movements were recorded in a variety of conditions that are known to elicit stereotypical and well-characterized oculomotor behaviors.

### Performance with artificial eyes

We first examined the noise of the system in the absence of any movements of the eye. As pointed out before, the human eyes are always in motion, even when attempting to maintain steady fixation on a point. Thus, an artificial eye is necessary to evaluate stationary steadystate performance. To this end, we used an artificial eye from Ocular Instruments (OEMI-7 model), which is designed to faithfully replicate the size, shape, and refraction characteristics of the human eye. This model eye includes an external transparent interface replicating the cornea, anterior and posterior chambers filled with 60% glycerine-water solutions^22^, and a lens made of polymethyl methacrylate that mimics the crystalline lens of the human eye.

We mounted the model eye on a dovetail rail attached to the eye-tracker’s stage so to place it at the position where the subject’s eye is normally at rest, aligned with the apparatus optical axis. We then tuned the eye-tracker to obtain clear and sharp Purkinje images, as we would do with a human eye. The eye-tracker’s output signal was recorded as the artificial eye remain immobile in this position. The variability of this signal sets a limit to the precision— *i.e.*, the repeatability of a measurement—that can be achieved by the system and provides information on its resolution, the minimum rotation that can be reliably detected. The data in Fig. 6A show that the noise of the system is within 0.1’, allowing in principle measurement of extremely small displacements.

**Figure 6:**
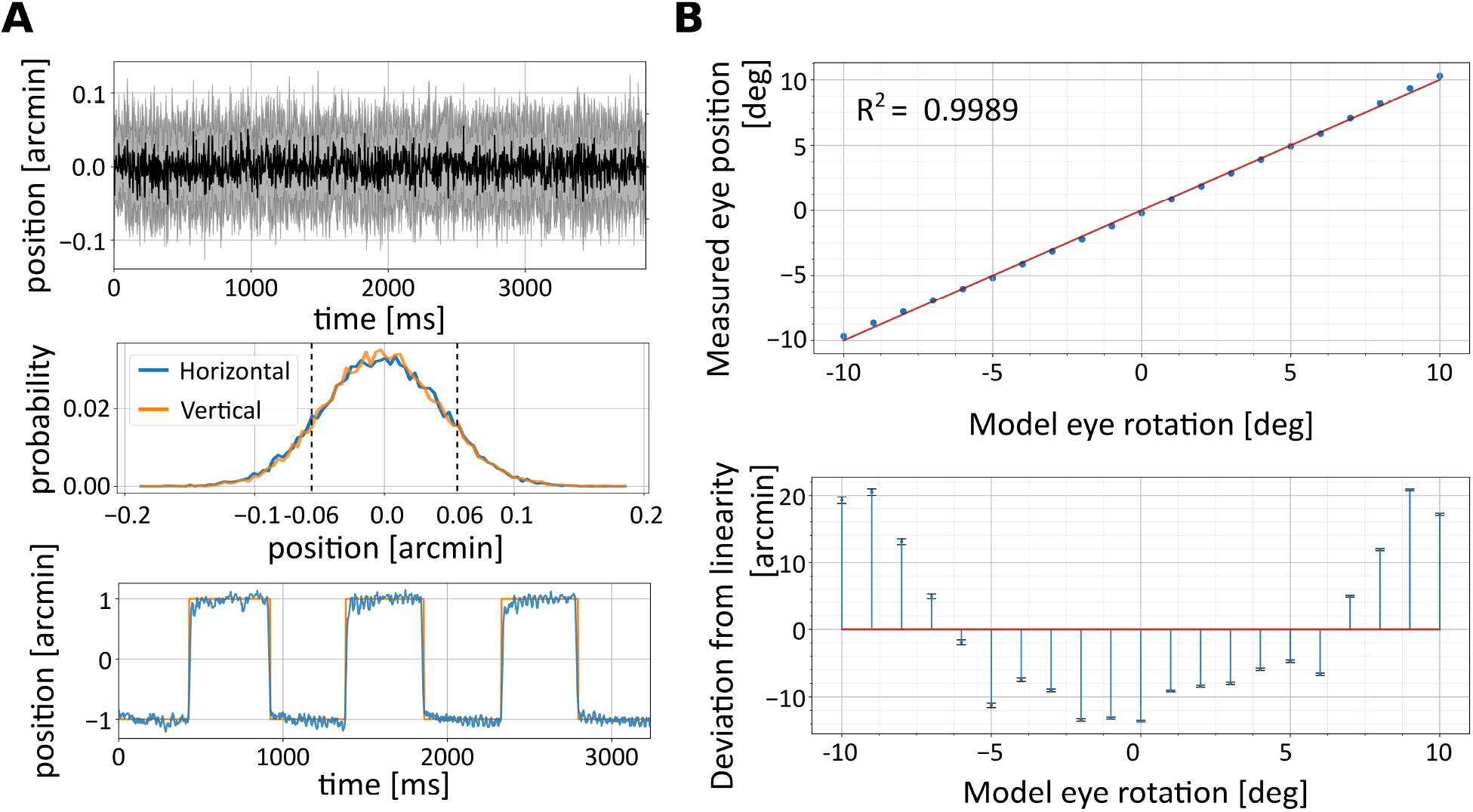
Performance with artificial eyes. (**A**) dDPI resolution. (Top) Average dDPI output (black line) ± one standard deviation (shaded region; N=30) measured with a stationary eye. (Middle) The resulting output distributions on both horizontal and vertical axes. Dashed lines represent one standard deviation. (Bottom) Measurements obtained when the eye moves following a 1’-amplitude square wave at 1 Hz. The orange line represents the ground truth, as measured by the encoders of the galvanometer controlling the eye. (**B**) dDPI linearity. (Top) Output measured when the artificial eye was rotated over a 20° range. The red line is the linear regression of the data. (Bottom) Deviation from linearity in the measured eye position.

We then used the artificial eye to examine performance under controlled rotations. To confirm that the system is indeed capable of resolving very small eye movements, we placed the eye at the same position as before, but now mounted on a custom stage controlled by a galvanometer (GSI MG325DT) rather than on the stationary rail. This apparatus had been previously calibrated by means of a laser to map driving voltages into very previse measurements of displacements, enabling rotations of the model eye with sub-arcminute precision. The bottom panel in Fig. 6*A* shows the response of the eye-tracker when the artificial eye was rotated by a 1-Hz square wave with amplitude of only 1’. Measurements from the eyetracker closely followed the displacements recorded by the galvo encoders, showing that that the system possesses both excellent resolution and sensitivity.

Our calibrated galvanometer could only rotate the artificial eye over a relatively small amplitude range. To test performance over a wider range, we mounted the artificial eye on a manual rotation stage (Thorlabs PR01/M) and systematically varied its orientation in 4° steps over 20°. The data points in Fig. 6*B* show that the eye-tracker correctly measures rotations over the entire range tested. In fact, as expected from the ray-tracing simulations, the system operates almost linearly in this range, with deviations from linearity smaller than 2% (20’; Fig. 6*B*). During tracking of human eyes, these deviations are further compensated by calibrations. But these data show that the need for non-linear compensations are minimal, at least with an artificial eye.

### Performance with human eyes

Given the positive outcome of the tests with the artificial eye, we then examined performance with human eyes. Thirty-three subjects (21 females, 12 males, age 24±5 years) participated in a variety of tests. Subjects provided their informed consent and were compensated according to the protocols approved by the University of Rochester Institutional Review Board.

As it is customary with most eye-trackers, before collecting data, preliminary steps were taken to ensure optimal tracking. These included positioning the subject in the apparatus and fine-tuning the offsets and focal length of the imaging system to obtain sharp, centered images of the eye. As normally done with the analog DPI, head movements were minimized by using a a dental bitebar and a headrest. Subjects either observed the stimulus directly, or, if they were not emmetropic, through a Badal optometer that corrected for their refraction. This device was placed immediately after the dichroic mirror and did not interfere with eyetracking. A preliminary calibration enabled conversion of the dDPI’s raw output in units of camera pixels into visual angle. To do this, we applied the same two-step gaze-contingent procedure that we have used previously to map the voltage signal from the analog DPI into visual angles^6^. This method relies on feedback from the subject to tune a bilinear calibration and has been shown to yield accurate gaze localization^5^.

In the first set of tests, we examined the robustness of the system to saccades, the abrupt, high-speed gaze shifts that occur frequently during natural tasks (Fig. 7A). We recorded saccades as observers freely viewed natural images, searched for targets in noise fields^11,23^, and executed cued saccades^24^. These tasks elicited saccades of all possible sizes, with amplitudes reaching over 12° and peak speeds up to 400°/s. The dDPI exhibited no difficulty in tracking saccades, yielding traces that were qualitatively indistinguishable from those recorded with the analog device (see example in Fig. 7A). The measured saccade dynamics exhibited the stereotypical linear relationship between saccade peak speed, duration, and saccade amplitude known as the main sequence^25^. These characteristics were virtually identical to those previously recorded with the analog device^26^ (*cf.* blue and orange lines in Fig. 7B).

**Figure 7:**
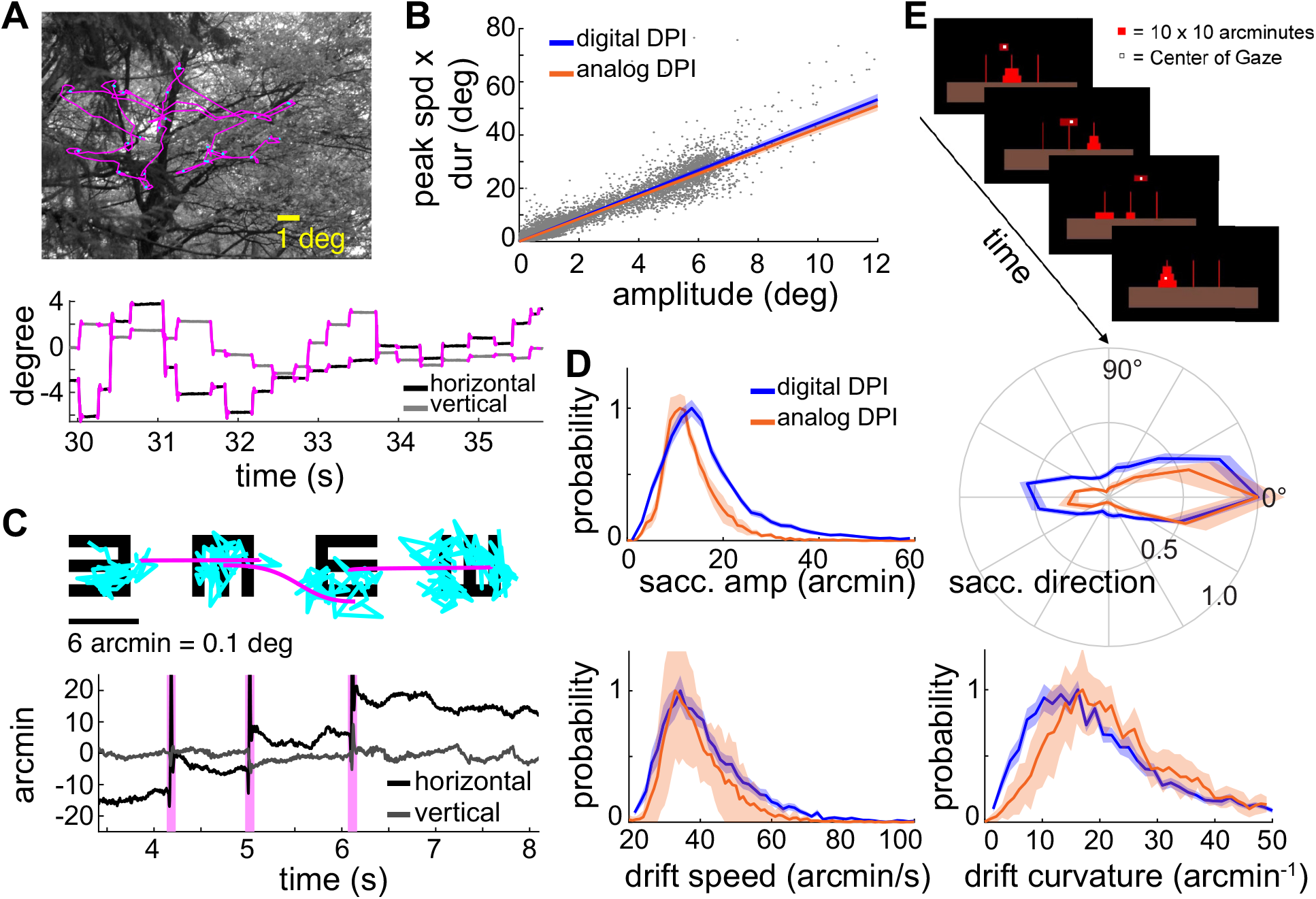
Human eye movements recorded by the dDPI. (**A**) Example of eye move-ments during exploration of a natural scene. An eye trace is shown both superimposed on the image (top) and displayed as a function of time (bottom). (**B**) Saccade main sequence. Data were collected from *N* = 14 observers during directed saccades or free viewing of natural images. The orange line represents the regression line of the main sequence for saccades measured in an analog DPI ^26^. (**C-D**) Eye movements recorded during examination of the 20/20 line of an acuity eye-chart. (*C*) An example of oculomotor trace displayed as in A. Microsaccades and drifts are marked in pink and blue respectively. (*D*) Characteristics of fixational eye movements. Data represent distributions of saccade amplitude, saccade direction, ocular drift speed, and drift curvature averaged across *N* = 26 subjects. Data are very similar to those recorded by means of the analog DPI. Lines and shaded errors represent averages and SEM across observers. (**E**) Gaze-contingent control of a miniaturized version of the Tower of Hanoi game. All participants were able to effortlessly pick up discs by fixating on them, move them by shifting their gaze, and drop them by blinking. Very accurate gaze localization is necessary for this task, since the thickness of each disk was only 10’.

We next examined the performance of the dDPI when observers executed small eye movements. Previous studies with search coil and analog DPI systems have extensively characterized how the eyes move during periods of fixation, when occasional small saccades (microsaccades;^27–29^) separate periods of slower motion (ocular drift^30–32^). We have recently shown that both microsaccades and ocular drift are controlled in high-acuity tasks ^6,8,33,34^, often yielding highly predictable behaviors. A particularly clear example is given by examination of the 20/20 line of an eye chart, a task in which microsaccades systematically shift gaze from one optotype to the next, and the drifts that occur on each optotype are smaller than those present during maintained fixation on a marker^9^. Given the robustness and repeatability of these effects, we repeated this task using the dDPI. Observers (*N* = 26) were instructed to report the orientations of tumbling Es in the 20/20 line of a Snellen eye chart. Optotypes were separated by just 10-12’ center to center, and each optotype only spanned 5-6’.

All subjects exhibited the typical oculomotor pattern previously reported^9^ with microsaccades systematically shifting gaze rightwards from one optotype to the next (pink curves) and drifts focused on each optotype (blue points, Fig. 7C). The characteristics of microsaccades and ocular drift measured by the dDPI were almost identical to those measured with the analog DPI (top panels in Fig. 7D). Ocular drifts recorded with the digital and analog devices also exhibited similar distributions of speed and curvature (bottom panels in Fig. 7D). These results corroborate the observations made with artificial eyes and show that the dDPI has sufficient temporal and spatial resolution to capture the characteristics of fixational eye movements.

We also tested the accuracy of the system—the capability of localizing gaze in the scene— by implementing a task that could only be carried out with accuracy better than a few arcminutes (Fig. 7E). In this task, observers performed a miniature version of the Tower of Hanoi game, a game in which disks need to be piled on poles. However, rather than moving disks with a mouse, participants used their gaze. They picked an object by fixating on it, moved it by shifting their gaze, and dropped it at a desired location by looking there and blinking. Since the disks were only 10’ thick, execution of the task was only possible with gaze localization more accurate than this. We implemented the task in EyeRIS, our custom system for gaze-contingent display^4^ at a refresh rate of 240 Hz, so that the average delay between measurement of eye position and update of the display was less than 8 ms. All subjects were able to execute the task without any training. They learned to interact with the scene in a matter of a few seconds and were able to position objects at the desired locations reliably and effortlessly. Thus, the speed and accuracy afforded by the dDPI enables robust gaze-contingent control of the display.

## Discussion

In the past two decades, major advances in the development of camera sensors and digital image processing have led to the flourishing of video-based eye-trackers. These systems have extended the use of eye-tracking far beyond laboratory use, enabling broad ranges of applications. However, most systems fall behind the resolution and accuracy achieved by the most sophisticated traditional approaches, so that the resulting performance is not yet at the level required by all vision science studies. In this article, we have described a digital implementation of the Dual Purkinje Images approach, a method originally developed by Crane and Steele that delivers oculomotor measurements with high temporal and spatial resolution. Our prototype shows that imaging technology is now sufficiently mature for digital video-oculography to replace the traditional gold-standard tools of oculomotor research.

Several factors contribute to the need for eye-trackers that possess both high precision and accuracy. A strong push for precision comes from the growing understanding of the importance of small eye movements for human visual functions. The last decade has seen a surge in the interest for how humans move their eyes at fixation, the very periods in which visual information is acquired and processed, and considerable effort has been dedicated to studying the characteristics and functions of these movements and the resulting motion on the retina. Precision and sensitivity are needed in these studies to reliably resolve the smallest movements. Furthermore, when combined with minimal delay, high resolution also opens the door for gaze-contingent display control, a powerful experimental tool that enables control of retinal stimulation during normal oculomotor behavior (Santini et al, 2007). This approach allows rigorous assessment of the consequences of the visual flow impinging onto the retina, as well as decoupling visual contingencies and eye movements.

In terms of accuracy, the capability of finely localizing the line of sight in the visual scene is critically important not just in vision science, but also in a variety of applications. Multiple studies have shown that vision is not uniform within the foveola ^11,35,36^ and humans tend to fixate using a very specific sub-region within this area (the preferred retinal locus of fixation), a locus which appears consistently stable across tasks and stimuli^8,9,37,38^. Accurate localization of the point in the scene projecting to this locus is essential in research questions that involve precise absolute positioning of stimuli (*e.g.,* presentation of stimuli at desired eccentricities), as well as in applications that require fine visuomotor interaction with the scene (*e.g.,* using the eyes as a mouse to interact with a computer).

Unfortunately gaze localization represents an insidious challenge, as video-based eye-trackers do not directly measure gaze position in space, but need to convert their raw measurements in visual coordinates, an operation that adds uncertainty. The parameters of this conversion are usually estimated via calibrations with observers fixating at known locations. However, both inaccuracies in the data points themselves and interpolation errors, particularly in highly non-linear eye-trackers, contribute to lower accuracy. As mentioned before, in normal, untrained, observers, just the error caused by eye movements during fixation on each of the points of a calibration grid can be as large as the entire foveola ^39^. In keeping with these considerations, accuracy is reported to be around 0.3-1° for video eye-trackers^40,41^, values that likely under-estimate the actual errors.

As demonstrated empirically by the real-time scene interaction described in Fig. 7E, gaze localization in our apparatus is sufficiently accurate to allow observers to use their gaze to effortlessly pick and drop objects as small as 10’. This high degree of accuracy is primarily the consequence of two factors. A major contribution comes from the specific, monotonic relation between displacements in the Purkinje images and angle of eye rotation, particularly in the central region where the mapping is almost linear (Fig. 4D;^13^). This linearity provides the foundation for the accuracy of the analog DPI (around 0.1°^42^ in trained observers) and is little affected by the physiological individual variations in anatomical and optical characteristics present in healthy subjects (see Fig. 2B). The second important component is the conversion of precision into accuracy resulting from gaze-contingent calibration^4^, a procedure in which subjects iteratively refine the mapping on the basis of real-time visual feedback, effectively improving accuracy by almost one order of magnitude^5,6^.

The combination of resolution, accuracy, and real-time gaze-contingent control afforded by the analog DPI has been extensively used to study visual functions in the presence of eye movements. In our laboratory, these techniques have been applied, among other things, to examine the influence of eye movements on spatial vision^43,44^, map visual functions across the fovea^11^, study the interaction between eye movements and attention^10,45^, and elucidate oculomotor strategies in both microsaccades^7,20^ and ocular drifts^46^. For example, accurate gaze localization has been instrumental to unveil that microsaccades precisely position the center of gaze on relevant features of the stimulus^6,7^ and that eye movements enhance fine spatial vision^9,47^. These studies were possible thanks to the capability of reliably resolving eye movements at arcminute level, accurately localizing gaze, and modifying stimuli in realtime, all capabilities that are now afforded by the system described here.

The dDPI presents several advantages over its analog predecessor. Although very precise, the analog DPI is cumbersome both to use and maintain. Maintenance requires frequent tuning for optimal performance, and the experimenter needs to have good technical knowledge of the apparatus to operate it effectively. In contrast, the dDPI is easy to operate and requires little maintenance, like most video-based eye-trackers^48^. It achieves higher stability than its analog predecessor, as it does not rely on the mechanical motion of detectors. And it can likely achieve higher resolution as well, as the estimation of the Purkinje’s positions is based on many pixels, rather than just four quadrants as in the analog system.

A further advantage of the dDPI is that it can be flexibly configured to the needs of the study under consideration. Resolution can be easily traded for width of the field of view, by modifying the optics. For example, in studies that focus on small eye movements, narrowing the imaged portion of the eye yields more pixels in the regions of interest, and therefore higher resolution. Flexible control of this trade-off can be achieved with a zoom lens. Similarly, temporal resolution can be flexibly modified and traded with spatial resolution by varying the size of the output image or changing the camera. In fact, the dDPI allows higher temporal acquisition rates than those afforded by the analog DPI, which is limited by its internal electrical and mechanical filtering. The processing algorithm can also be substantially improved. In our proof-of-concept prototype, we emulated the tracking algorithm followed by the analog machine. But the flexibility and intelligence afforded by software programming enables the development of more robust algorithms. This is particularly true if time is not of essence, and eye movements can be processed offline.

In sum, our dDPI allows high-resolution, real-time eye-tracking while circumventing many of the difficulties associated with traditional high-precision devices, such as the analog DPI and eye coils. The resulting robustness and operational simplicity may bring high-resolution eyetracking outside the realm of specialized laboratories. The system’s flexible and customizable design should facilitate integration with devices that require fine oculomotor measurements and/or compensation for eye movements, such as retinal imaging devices. Furthermore, this development may open the door for studies in group of subjects that could not easily tolerate the constraints imposed by previous systems, such children, the elderly, and patients.

## Acknowledgements

This work was supported by Reality Labs and by NIH grant R01EY18363. We thank Martina Poletti, Soma Mizobuchi and the other members of the Active Perception Laboratory for helpful comments, discussions, and technical assistance during the course of this work.

